# A high quality reference genome for the fish pathogen *Streptococcus iniae*

**DOI:** 10.1101/2019.12.17.880476

**Authors:** Areej S. Alsheikh-Hussain, Nouri L. Ben Zakour, Brian M. Forde, Oleksandra Silayeva, Andrew C. Barnes, Scott A. Beatson

## Abstract

Fish mortality caused by *Streptococcus iniae* is a major economic problem in fish aquaculture in warm and temperate regions globally. There is also risk of zoonotic infection by *S. iniae* through handling of contaminated fish. In this study, we present the complete genome sequence of *S. iniae* strain QMA0248, isolated from farmed barramundi in South Australia. The 2.12 Mb genome of *S. iniae* QMA0248 carries a 32 Kb prophage, a 12 Kb genomic island, and 92 discrete insertion sequence (IS) elements. These include 9 novel IS types that belong mostly to the IS*3* family. Comparative and phylogenetic analysis between *S. iniae* QMA0248 and publicly available complete *S. iniae* genomes revealed discrepancies that are likely due to misassembly in the genomes of isolates ISET0901 and ISNO. We also determined by long-range PCR that a tandem duplication of an rRNA region in the PacBio assembly of QMA0248 was an assembly error. A similar rRNA duplication in the PacBio genome of *S. iniae* 89353 may also be a misassembly. Our study not only highlights assembly problems in existing genomes, but provides a high quality reference genome for *S. iniae* QMA0248, including manually curated mobile genetic elements, that will assist future *S. iniae* comparative genomic and evolutionary studies.

## Introduction

*Streptococcus iniae* is a fish pathogen that causes mortality in a wide range of fish species in wild and farmed, marine and freshwater environments, resulting in large economic losses to aquaculture (1, 2). *S. iniae* is also considered an opportunistic human pathogen, causing sporadic infections mostly in the elderly who have more than one underlying health condition such as diabetes mellitus, or chronic rheumatic heart disease (3, 4). *S. iniae* pathogenesis is imparted through a repertoire of virulence factors (VF) including surface proteins, secreted toxins, and capsular polysaccharide (CPS) (4). VFs can be acquired through lateral gene transfer (LGT) of mobile genetic elements (MGE) such as composite transposons, genomic islands (GI) or prophages.

MGEs are a means by which bacterial pathogens acquire traits that help adapt to changing conditions including vaccination, antibiotics, a new host or environment (5, 6). Indeed, they are considered the main drivers of gene flux in bacteria, contributing to diversity within species (7). Insertion sequence (IS) elements, for instance, are small MGEs (0.7-3.5 Kb) that have an important role in evolution and genome plasticity. IS insertion within bacterial chromosomes or plasmids can result in genetic modifications through insertional inactivation of genes or up-regulation of adjacent intact genes through outward-facing promoter sequences carried by some IS (8, 9). In some cases pairs of IS can mobilize intervening sequence as a composite transposon (8). The mobility of IS elements leads to their expansion or loss within bacterial lineages. Expansion is associated with accumulation of pseudogenes, which is considered an early stage in genome reduction as a mechanism for adaptation (9). Accordingly, to obtain the complete evolutionary picture within bacterial species it is important to study the distribution and abundance of IS elements.

As yet there is no study that focuses on the diversity and distribution of MGEs in *S. iniae* genomes. In fact, only four complete *S. iniae* genomes were deposited in Genbank at the commencement of the present study, none of which had comprehensive annotations for MGEs: SF1, YSFST01-82, ISET0901, and ISNO. *S. iniae* SF1 (accession: CP005941) was cultured from moribund flounder from a fish farm that experienced an epidemic in North China (10). YSFST01-82 (accession: CP010783) is an isolate from a diseased Olive flounder from South Korea (2012) (11). ISET0901 (accession: CP007586) is reported to be a highly virulent strain, which was isolated from diseased Nile tilapia during an outbreak in Israel (2005) (12). The strain ISNO (accession: CP007587) was obtained through selection for resistance of *S. iniae* ISET0901 to novobiocin (13).

There are currently 25 different IS families in the ISFinder database, but only one IS described for *S. iniae* (IS*Stin1* of the family IS*256*). As is typical for complete bacterial genomes, the annotation of the four complete *S. iniae* genomes available at Genbank are limited to some of the transposase genes associated with IS elements without definition of the IS boundaries. Indeed, the difficulty in annotating IS elements and the lack of reliable automated annotation methods means that only a small subset of complete genomes have accurate IS annotations. Small partial IS elements are often disregarded although they reflect valuable insights on ancestral recombination events. With the increasing availability of long-read sequencing there is a need for a high quality, well-characterized *S. iniae* reference genome that better enables the impact of IS elements on evolution and diversity to be determined.

In this study, we have completely characterized the genome of *S. iniae* QMA0248. Manual curation of annotations for IS, genomic islands, prophages, and CRISPR was carried out along with a comparison with the four publicly available complete genomes from NCBI (SF1, YSFST01-82, ISET0901, and ISNO). Comparative and phylogenetic analyses revealed discrepancies between the MGE content indicating likely misassembly in the genomes ISNO, and ISET0901. The complete genome of QMA0248 will provide an important scaffold for future phylogenomic studies of the *S. iniae* species.

## Results and Discussion

### Genomic features of *S. iniae* QMA0248

The genome of *S. iniae* QMA0248 consists of a single circular chromosome of 2,116,570 bp with no plasmids (Figure 1 and Table 1). The QMA0248 chromosome has an average GC content of 36.8 %, consistent with the other four *S. iniae* genomes (SF1, YSFST01-82, ISET0901, and ISNO) and in common with several other *Streptococcus* spp. such as *S. agalactiae* (35.6%) (14), *S. pneumoniae* (39.7%) (15), and *S. pyogenes* (38.5%) (16). Of note, there is a high degree of strand bias in the genome of *S. iniae* QMA0248 where genes are preferentially oriented in the leading strand, which is typical for Firmicutes (Figure 1). Only seven percent of QMA0248 protein-coding genes are annotated as hypothetical proteins compared with a quarter in SF1 and 11-14% in the other three published genomes: YSFST01-82, ISET0901, and ISNO. There are 68 pseudogenes identified in QMA0248 (GenBank assembly accession: CP022392.1), most due to interruption by IS (Figure 1, Supplementary Table S1). This is approximately five times greater than the number of pseudogenes predicted in SF1. Collectively, these differences between the compared genomes QMA0248, SF1, YSFST01-82, ISET0901, and ISNO are likely to reflect different approaches to annotation. The other 25 pseudogenes are caused by in-frame stop codons or frame-shifts, all supported by additional mapping of Illumina reads of the strain QMA0248 against its PacBio assembly.

**Figure 1:**
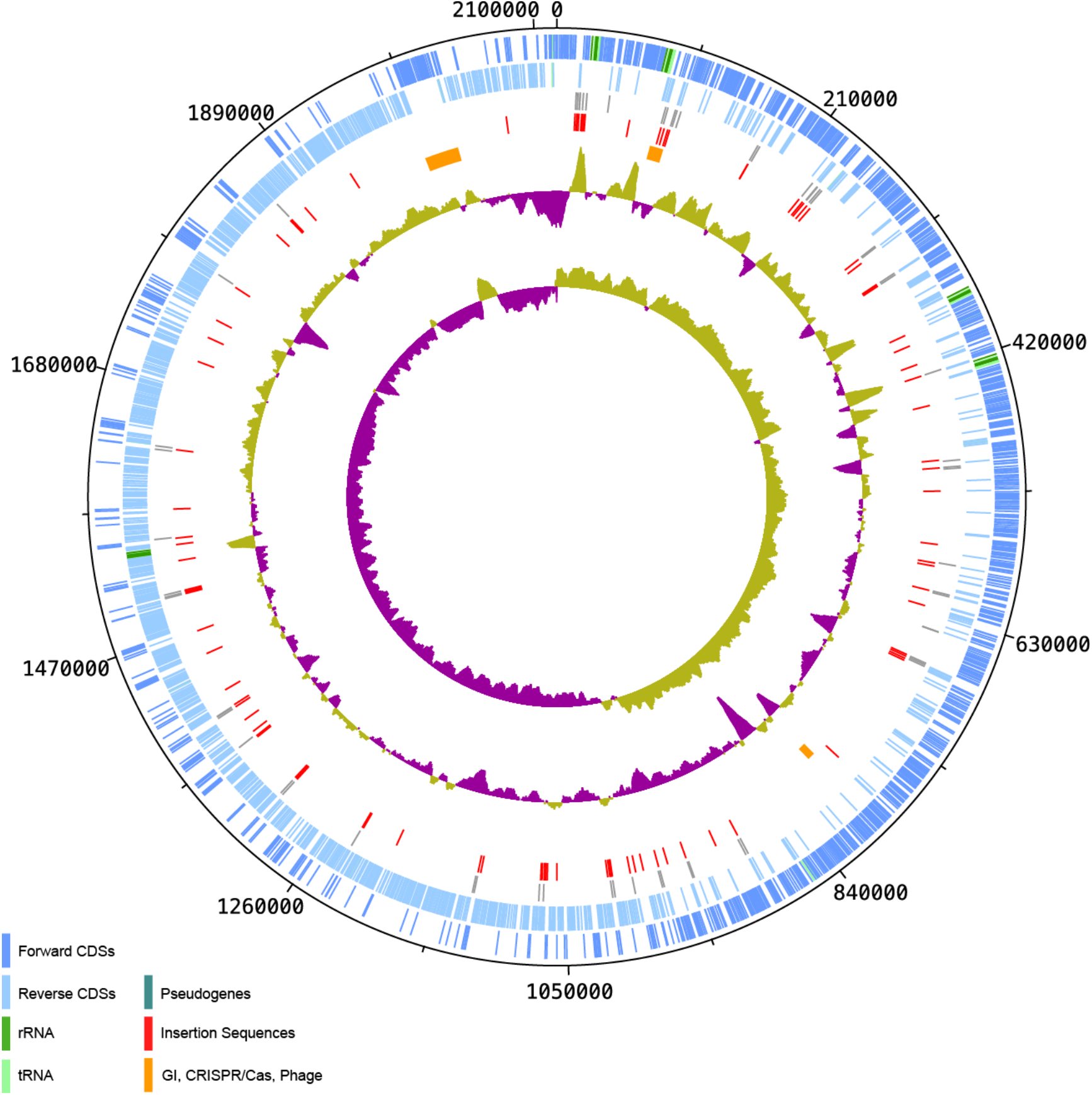
Circular map of the genome *Streptococcus iniae* QMA0248. Genomic features from outer ring to the inner ring are described in the key to the left, where the innermost two rings correspond to the GC skew (inner), and GC plot (outer). CDS: coding sequence. GI: genomic island. The circular map was generated using DNAPlotter (59).

**Table 1:**
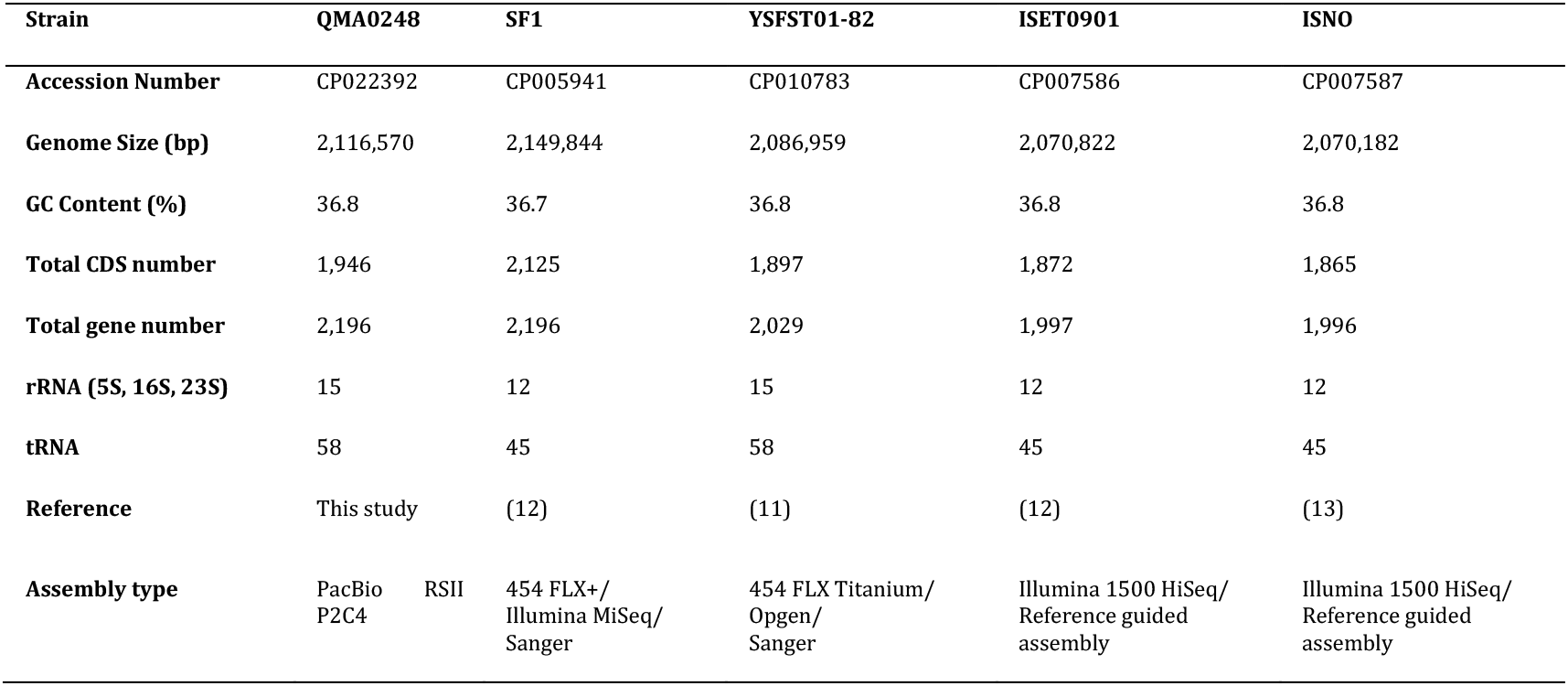
General features of five *S. iniae* complete genomes.

QMA0248 has a single CRISPR locus (Clustered regularly interspaced short palindromic repeats) (Figure 1), which harbors a tandem array of 15 identical 36 bp repeats, separated by 14 distinct 30 bp spacers, which is about double the size of the CRISPR array in SF1, ISET0901, and ISNO (Supplementary Figure S1). Upstream of QMA0248 CRISPR are four Cas genes, *csn2*, *cas2*, *cas1*, and *cas9*, which alongside the CRISPR locus provide adaptive immunity against foreign DNA (e.g. phage and plasmids) (17).

QMA0248 harbors 58 tRNA genes and 15 rRNA loci, consisting of 5S, 16S, and 23S genes, arranged in five loci. In contrast, there is one rRNA operon fewer in SF1, ISET0901, and ISNO than in QMA0248 (Table 1). Furthermore, during the preparation of this manuscript, a PacBio complete genome was published for the strain *S. iniae* 89353 (accession: CP017952) which has an identical rRNA arrangement to QMA0248 except that one rRNA locus encodes a ~7 Kb tandem duplication (i.e. six loci in total) (18). Such intra-species variation in the number of rRNA genes (and tRNA genes) is not uncommon in bacteria (including streptococci) (19, 20), and prompted us to investigate further.

### Confirmation of rRNA assembly in *S. iniae* QMA0248 genome

PacBio reads for QMA0248 were initially assembled into a large ~2 Mb contig representing most of the chromosome of *S. iniae* QMA0248 and three contigs less than 10 Kb in length. The short contigs appeared to be single read chimeras that were discarded from the final assembly. However, the identification of the tandem rRNA region in *S. iniae* 89353 prompted us to review the assembled short contigs. One of these ~7 Kb discarded contigs encoded an rRNA region (5S, 16S, and 23S genes in tandem with an intervening cluster of tRNA genes). Subsequent reassembly and visualization of mapped raw reads indicated that the additional rRNA contig could be placed in three of the five rRNA operon locations to form a ~13 Kb putative tandem duplication of 5S, 16S and 23 rRNA genes as seen in the 89353 genome (18). Closer examination of the read pileup for the ~7 Kb rRNA contig revealed that the tandem duplication was not well supported by overlapping reads (Supplementary Figure S2) suggesting that it may be a chimeric assembly of reads from more than one rRNA locus. To investigate the location of the putative tandem rRNA duplication in *S. iniae* QMA0248 we carried out long-range PCR across each of the five potential rRNA loci in the chromosome. PCR revealed no tandem rRNA duplication in any locus (Supplementary Figure S3). Never-the-less, we cannot rule out that during the culturing step, prior to DNA extraction and genome sequencing, there existed a sub-population of QMA0248 cells with the tandem rRNA duplication. This would be consistent with finding only a small number of reads spanning the tandem repeat. Similar doubt exists over the tandem rRNA repeat in 89353, which should also be investigated at the read-level.

### Characterization of large mobile genetic elements in *S. iniae* QMA0248

Our investigation of the rRNA discrepancies in *S. iniae* SF1, ISET0901, and ISNO also revealed differences in the mobile genetic element (MGE) content within the available complete genomes. The chromosome of QMA0248 has a single ~12 Kb genomic island (GI-Leu) inserted within the tRNA-Leu downstream of a large number of consecutive ribosomal genes (Table 2). GI-Leu encodes an integrase at its 5’ end (QMA0248_0125), which is predicted to be responsible for the GI insertion. A ~2.8 Kb region at the 5’ end of the GI (87881–90708) is homologous to the fish pathogen *S. parauberis* KCTC 11537, with 90% nucleotide sequence identity, including part of the integrase, plasmid replication gene, and two hypothetical proteins. The island encodes a collagen-binding surface protein (Cna, B-type domain), which is a virulence factor with an LPxTG cell wall anchor motif, a conserved Gram-positive *cocci* pentapeptide (21). An IS*30* family transposon insertion truncates the collagen-binding domain encoded by *cna*, likely leading to loss of functionality in QMA0248. This adhesin has been shown to play an important pathogenic role in *Staphylococcus aureus* by facilitating bacterial cells adherence to host collagen (22). Most of this GI appears to have been deleted in the genomes of SF1, ISET0901, and ISNO only, along with the rRNA operon upstream of it (Figure 3), which explains the difference with QMA0248 in total number of rRNA and tRNA genes (Table 1).

**Table 2:**
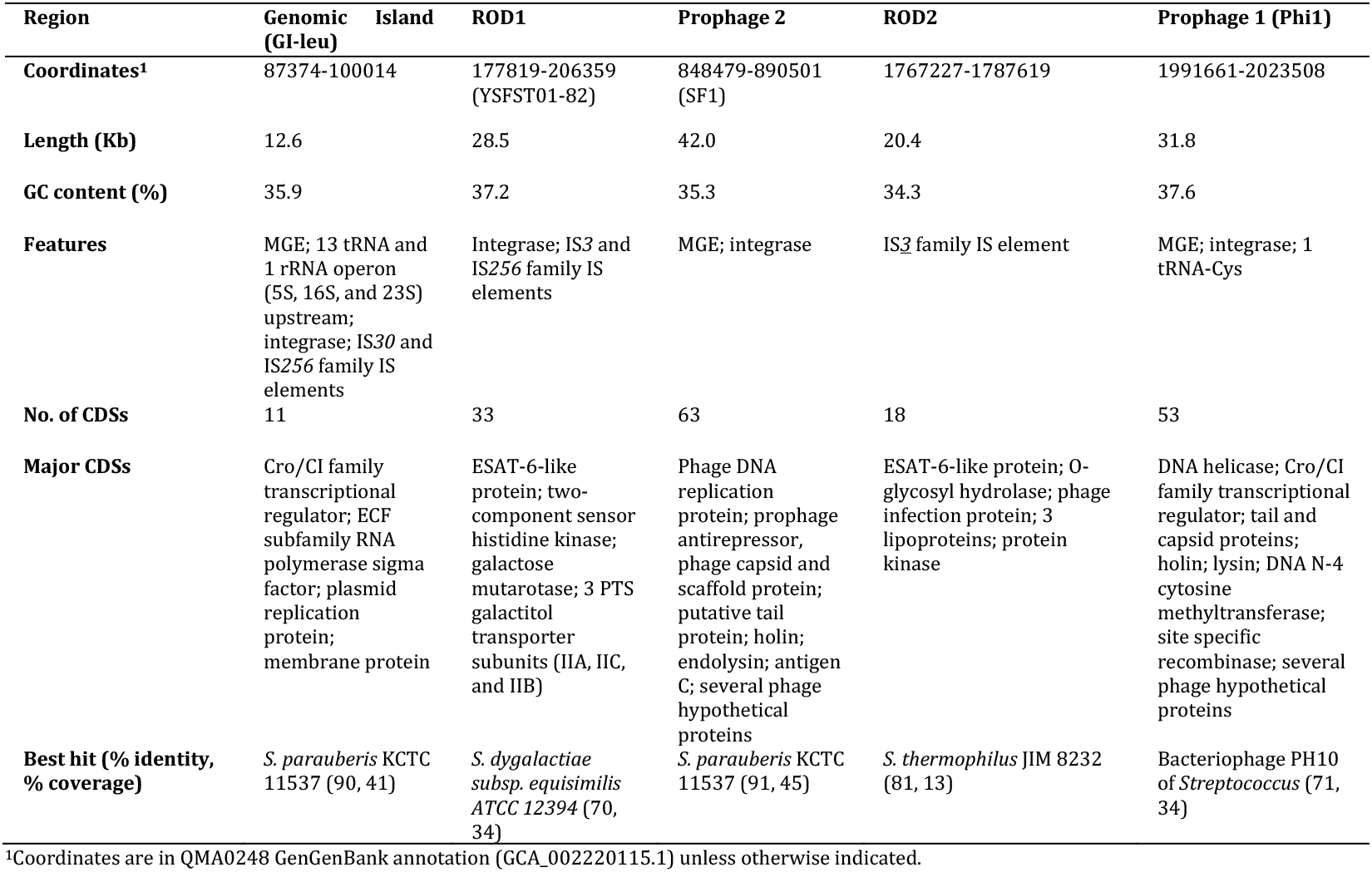
Large mobile genetic elements (MGE) and regions of difference (ROD) identified in the 5 *S. iniae* genomes analyzed (QMA0248, SF1, YSFST01-82, ISET0901, and ISNO).

The genome of *S. iniae* QMA0248 harbors a single ~32 Kb incomplete phage (Phi1) (1991661–2023508), including a 5’ integrase gene (QMA0248_1936), inserted upstream of a tRNA-Cys gene (Table 2). A total of 44 genes encoding phage proteins were identified, including genes involved in DNA replication such as DNA polymerase III, tail morphogenesis, as well as host lysis such as holin and lysin, in addition to 24 phage hypothetical proteins. More than half of the genes encoding phage proteins carried by QMA0248 are homologous to proteins encoded by temperate bacteriophage *Streptococcus* PH10 (56.8% according to PHAST) (23). Furthermore, Phi1 in QMA0248 exhibits a remarkable nucleotide sequence identity (99%) to a prophage encoded within the SF1 genome in the same locus, whereas it is entirely absent in YSFST01-82, ISET0901, and ISNO (Figure 3).

### Characterization of *S. iniae* QMA0248 insertion sequences

Insertion sequences (IS) were analyzed in the *S. iniae* QMA0248 genome using the ISFinder database coupled with manual curation. The analysis revealed 92 IS (Table 3), which is higher than the average number per bacterial genome (n=38) but consistent with the lifestyle of *S. iniae* as a facultative pathogen (24, 25). Furthermore, the number of IS found in *S. iniae* QMA0248 is substantially higher than other Streptococci such as *S. mitis* strain B6 (n=63) but comparable to that of the Gram positive fish pathogen *Lactococcus garvieae* (26, 27). The 92 IS elements belong to 7 different IS families and 20 IS types. These include 9 novel types belonging to IS*3*, IS*30*, IS*1182*, and IS*200*/IS*605* families, which we have submitted to the ISFinder database (IS*Stin2–*IS*Stin10*) (Table 3 and Supplementary Tables S2–4). Around half of all IS copies in QMA0248 belong to these 9 novel types consistent with expansion of *S. iniae* specific IS since speciation (Table 3). Amongst those genes disrupted by IS in QMA0248 is the restriction enzyme component of a type II Restriction Methylation system that probably recognizes “GCNGC” (28). This insertion renders the cognate methyltransferase (MTase) (QMA0248_0516) an orphan and, given the high number of GCNGC sites (3814/Mb in QMA0248) across the genome, suggests a potential role for this MTase in global gene regulation. By comparison, there are 7762/Mb available GATC sites in the *E. coli* K-12 MG1655 genome available for methylation by Dam, the archetypal orphan type II MTase known to play a major role in gene regulation (29). Further discussion of the methylome data generated by PacBio sequencing of QMA0248 is provided as Supplementary Information. . Taken altogether, the *S. iniae* genome harbors a large repertoire of IS elements, which may be associated with adaptation to host or environment. Indeed, it is well accepted that IS expansion is an early sign of genome reduction as a mechanism of adaptation to host (25, 30, 31).

**Table 3:**
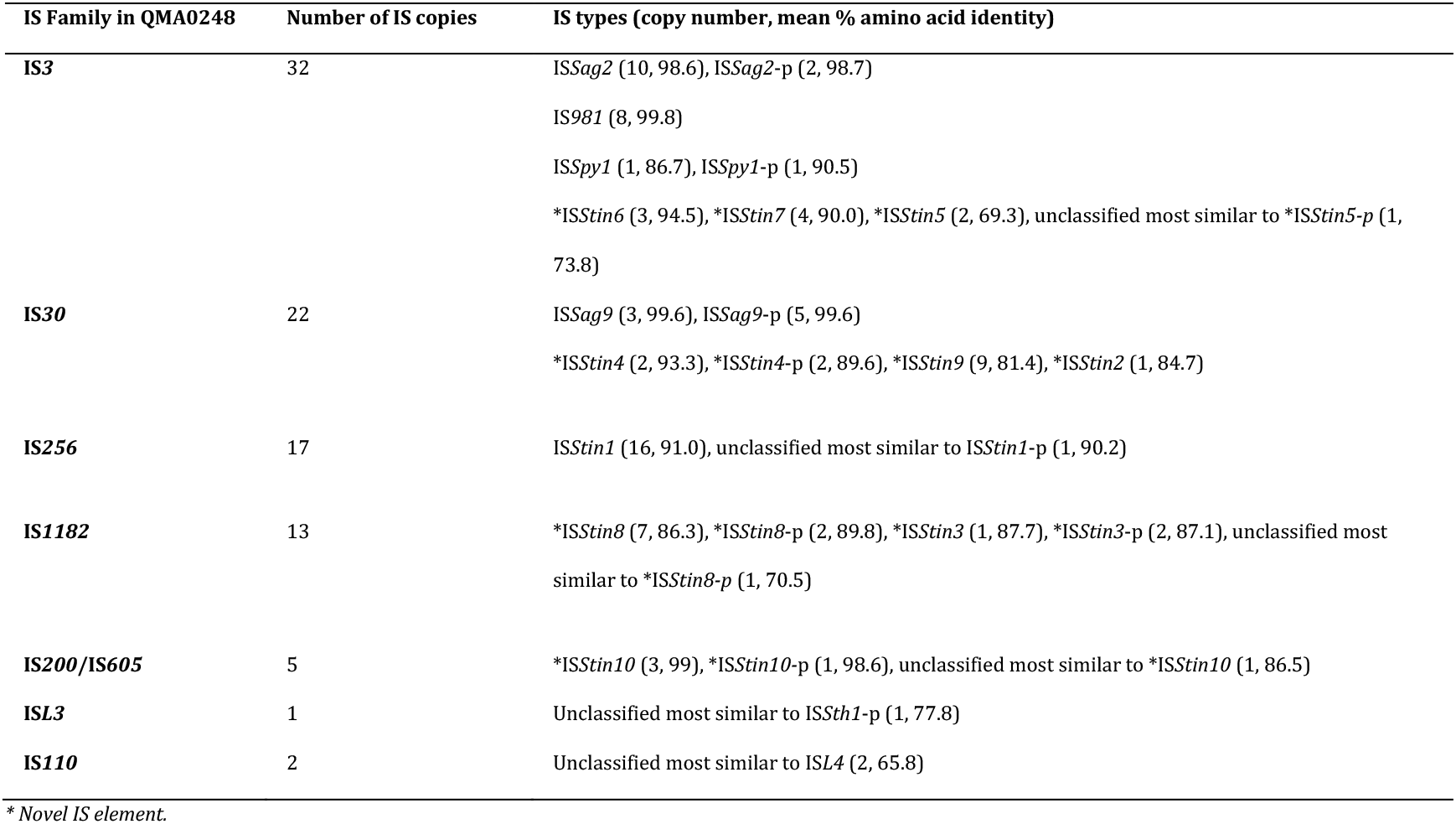
Summary of all insertion sequences (IS) identified in QMA0248. Partial IS are suffixed by −p.

### Phylogenetic and comparative analysis of *S. iniae* QMA0248, SF1, YSFST01-82, ISET0901, and ISNO

The core genome of *S. iniae* QMA0248, SF1, YSFST01-82, ISET0901, and ISNO accounts for ~75% of the chromosome. IS were compared between the reference chromosome of QMA0248 and each of the *S. iniae* chromosomes (SF1, YSFST01-82, ISET0901, and ISNO) using Artemis Comparison Tool (ACT) (32). Although IS elements typically result in genomic rearrangements and loss of synteny, this is not seen in *S. iniae.* This lack of rearrangement is reflected by the consistent pattern of GC skew in the genome of *S. iniae* QMA0248 (Figure 1). Eight IS copies out of the 92 detected in QMA0248 are absent in the genomes of SF1, ISET0901 and ISNO only (Supplementary Table S2). Other IS elements are unique to the genomes SF1, ISET0901 and ISNO in syntenic positions. An interesting example is an IS*981* (SF1 locus_tag: K710_0799 and K710_0800), inserted in *cas9* gene in the CRISPR/Cas region only in those three genomes (Figure 2). Another example is IS*Stin1*, inserted only in SF1, ISET0901, and ISNO (SF1 locus_tag: K710_0761). Additionally, three IS copies are present in syntenic positions only in QMA0248, ISET0901, and ISNO. Eight insertions are absent in YSFST01-82 only and another seven IS copies are found exclusive to QMA0248 (Supplementary Table S2).

**Figure 2:**
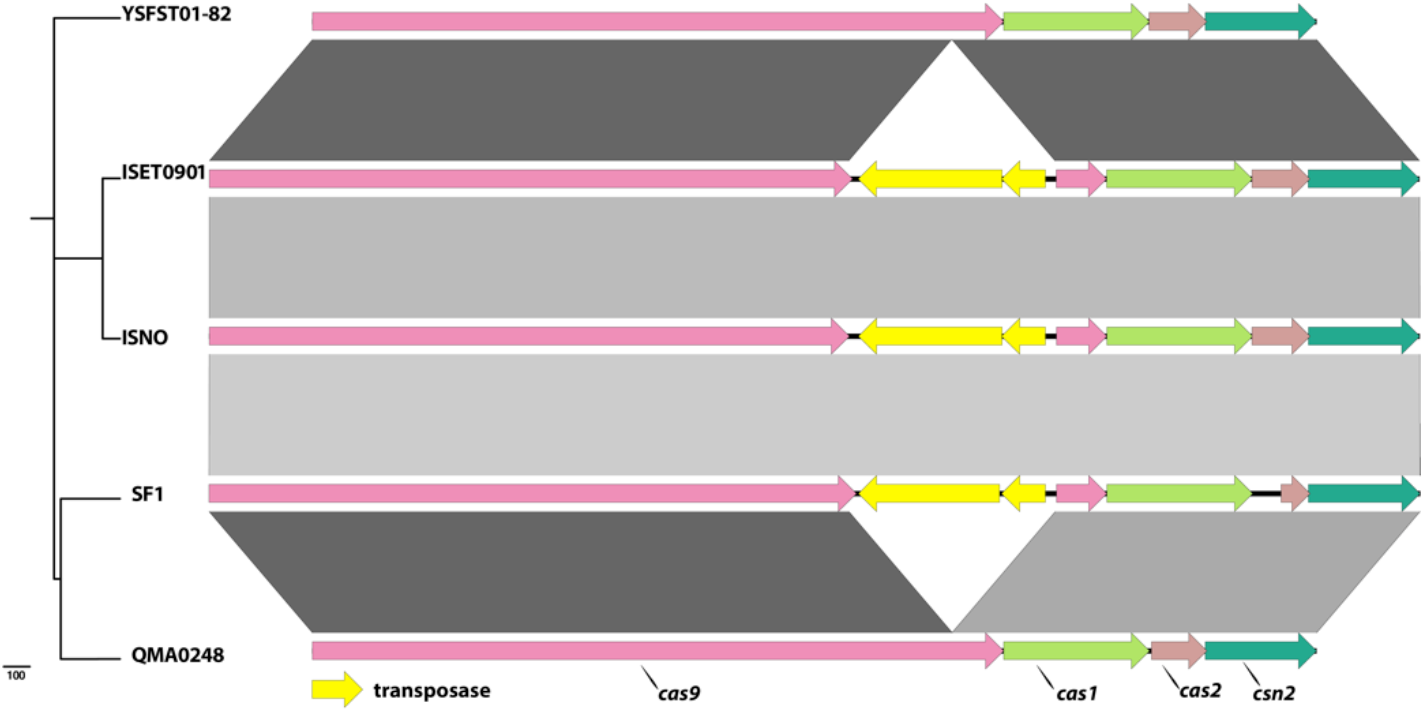
Comparison of the CRISPR/Cas region between QMA0248, SF1, YSFST01-82, ISET0901, and ISNO. Alignment of Cas genes where the genomes are ordered according to their position in the phylogenetic tree (left). The maximum Likelihood (ML) phylogeny was rooted to QMA0140 (not shown) and built using 1,111 SNPs. Arrows correspond to Cas genes, which are labeled at the bottom. Figure was produced using EasyFig (50) using 500 bp as minimum length, 90% as minimum identity value, and 0.001 as maximum e-value.

**Figure 3:**
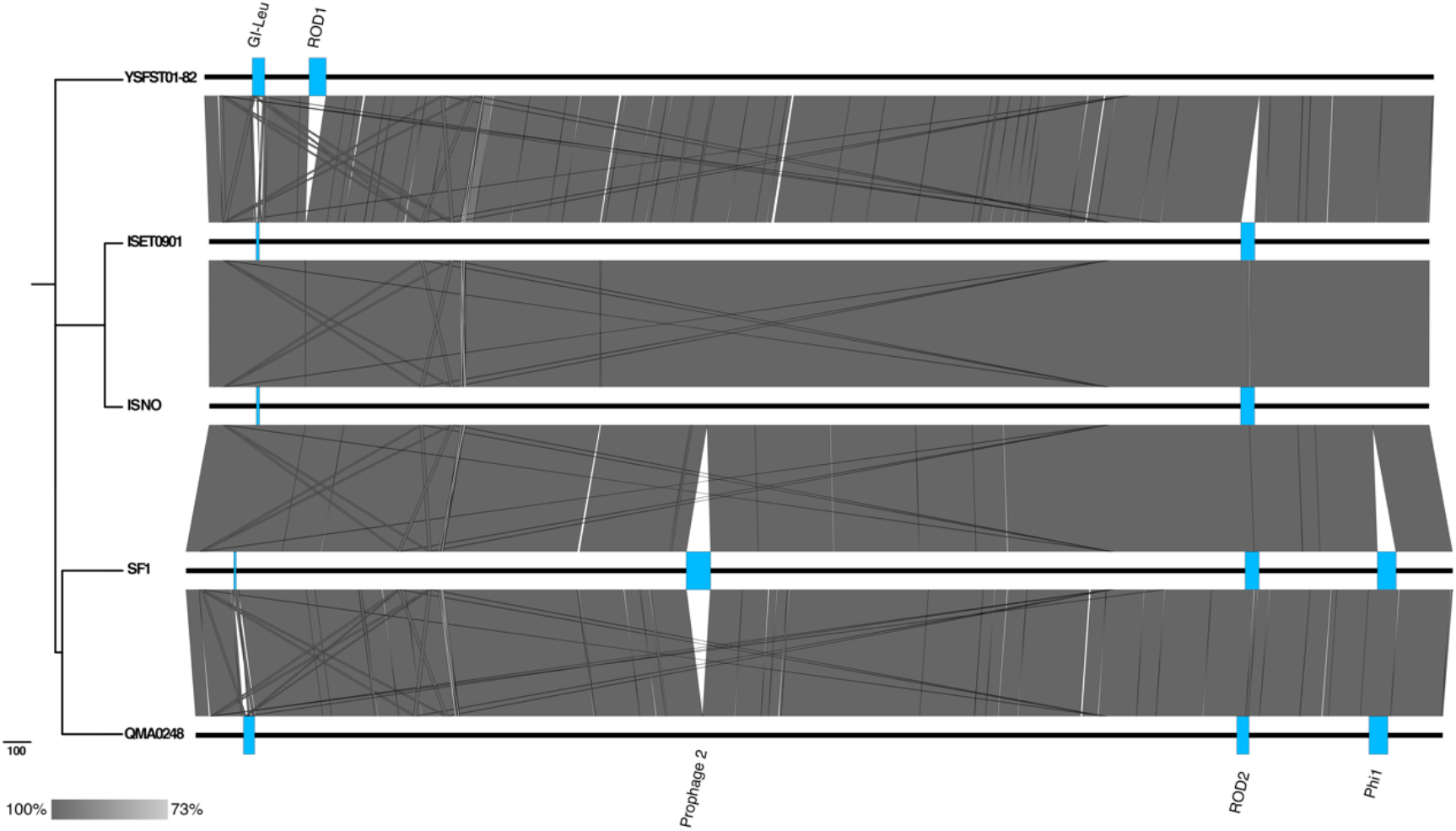
Whole-genome alignment of the 5 genomes QMA0248, SF1, YSFST01-82, ISET0901, and ISNO. The genomes are ordered according to their position in the core SNP-based phylogenetic tree. The maximum Likelihood (ML) phylogeny was rooted to QMA0140 (not shown) and built using 1,111 SNPs. The scale bar indicates the number of substitutions represented by branch lengths. BLASTn comparison was produced using EasyFig (50) using 2000 bp as minimum length, 50% as minimum identity value, and 1 × 10^−17^ as maximum e-value.

In addition to the IS differences, the five *S. iniae* genomes have major differences in the content of other mobile genetic elements (MGE) that reflect variations in the length of their respective chromosomes (Table 1). These MGEs include two prophages (Phi1 and Phi2), a genomic island (GI), and two other regions of difference (ROD) (Table 2, Figure 3). This includes a ~28 Kb region that is only found in YSFST01-82 (ROD1), and a ~20 Kb region that is present in four genomes but almost entirely absent from YSFST01-82 (ROD2) (Table 2, Figure 3).

Most variations in MGEs (including IS) were found to be incongruent with the core SNP phylogeny. For instance, the deleted ~12 Kb GI-Leu and the *cas9* gene disrupted by IS*981* exist only in SF1, ISET0901, and ISNO (Figures 2 and 3), but these three isolates appear on divergent branches, indicating potential independent events (Supplementary Figure S4). To investigate the discrepancies between MGEs and phylogeny we compared multiple phylogenetic trees that were constructed using different methods, including the core genome, core SNP and using different software (Supplementary Figure S4). All phylogenies consistently revealed that *S. iniae* isolates QMA0248 and SF1 cluster together in one clade, whereas ISET0901 and ISNO cluster in another, and all four isolates cluster separately to YSFST01-82, the latter diverging earliest from the root (Supplementary Figure S4).

### Discrepancies between the *S. iniae* genomes are likely due to misassembly

Taken together, our comparative analyses suggest that one or more of the *S. iniae* genomes under comparison have been misassembled. The genomes ISET0901 and ISNO were both assembled from Illumina data using BioNumerics (Applied Math) and the genome *S. iniae* SF1 as a reference (12, 13). *S. iniae* SF1 was assembled *de novo* from a combination of 454 GS FLX+, Illumina MiSeq and Sanger sequencing (12). During the preparation of the present manuscript SF1 was removed from the NCBI RefSeq database. YSFST01-82 was also a hybrid assembly (454 GS FLX Titanium, Opgen optical mapping and Sanger sequencing) but it remains in the RefSeq database and is the designated genome for *S. iniae* at NCBI (https://www.ncbi.nlm.nih.gov/genome/?term=streptococcus+iniae; accessed 23^rd^ Jan 2020).

We have no reason to suspect that the YSFST01-82 genome assembly is inferior to that of QMA0248, but adopting the latter as an alternative representative *S. iniae* genome is justified for investigators wishing to take advantage of a manually curated set of MGEs. In contrast we strongly recommend not using ISET0901 or ISNO in future comparative studies of *S. iniae* genomes. Reference-guided assembly was introduced to enable comparisons between two very closely related isolates. However, this practice can result in the erroneous inclusion of MGEs that exist in the template genome but are absent from the comparison strain. Even with careful curation it is impossible to avoid misplacing repetitive sequences such as IS, as observed here in the case of *cas9* insertion and the other 8 IS copies that are absent in SF1, ISET0901, and ISNO only. Moreover, reference-guided assemblies may result in the loss of novel regions that are only present in the newly sequenced strain, in which case a *de novo* approach is always required (33). Although reference-guided assembly is no longer generally accepted for prokaryote genomes, a number of examples remain available in public repositories such as GenBank. For both ISET0901 and ISNO the assembly strategy is clearly outlined in the comment field of the GenBank file, and the primary publications (12, 13). Never-the-less, the consequences of using such genomes in downstream analyses may not be apparent to all (for example, all three genomes are available in widely used genome databases such as PATRIC (www.patricbrc.org version 3.5.36)(34)).

Removal of some early hybrid 454 complete genomes from public repositories like RefSeq should help maintain the quality of available complete genomes. Long-read sequencing data from Pacific Biosciences and Oxford Nanopore bring complete bacterial genomes within reach of most laboratories, but here also significant care is often required to avoid misassembly. Furthermore, as illustrated here and in other studies (35, 36), what appear to be misassemblies may in fact be biologically relevant. Ultimately the onus is on the user of public data to exercise caution when validating the source, assembly strategy, and quality of available complete genomes.

## Conclusions

We assembled and annotated a high quality complete genome sequence for *S. iniae* QMA0248, including manual curation of 92 insertion sequences. Comparative analysis with publicly available complete genomes *S. iniae* SF1, YSFST01-82, ISET0901, and ISNO revealed discrepancies in the MGE content consistent with errors introduced by reference guided assembly of ISNO and ISET0901. Such problems are not new, but many bacterial genomes assembled in this way remain in public repositories of complete genomes. Our results emphasize the need to critically appraise complete genome assemblies prior to comparative analysis. Despite long-read sequencing becoming the gold standard for complete genome assembly of bacterial isolates, caution is needed to avoid misassembly. To better understand how insertion sequences, genomic islands, and other mobile elements contribute to *S. iniae* diversity, there is a need for larger genomic studies using global collections of *S. iniae* isolates from dissimilar origins. The genome of *S. iniae* QMA0248 represents an important resource for future *S. iniae* comparative genomic and evolutionary studies.

## Materials and Methods

### Bacterial strain and sequencing

*Streptococcus iniae* strain QMA0248 was isolated from diseased barramundi (*Lates calcarifer*) from a farm in South Australia in 2009 (37). Genomic DNA was prepared from several well-isolated colonies of *S. iniae* QMA0248 grown for 18 h on Todd-Hewitt agar from a master seed stock (non-subcultured) with the Genomic Tip 20 kit (Qiagen). Pre-incubation for 2 h at 37°C of cells suspended in 500 μL 50 mM EDTA containing 200 units of mutanolysin and 2 mg lysozyme was found to improve cell lysis prior to following the manufacturers protocol for purification of high molecular weight DNA. The genome of the strain QMA0248 was sequenced using 3 SMRT cells on the Pacific Biosciences (PacBio) RS II platform and P4C2 sequencing chemistry, which generated a total of 57,083 reads with an average length of 6,178 bp. Reads were deposited at BioProject <u>PRJNA385746</u> under accession <u>SRP109617</u>. The genome of the strain QMA0248 was also sequenced using Illumina Nextera libraries on HiSeq2000.

### Genome assembly and modified bases detection

PacBio sequencing reads derived from *S. iniae* QMA0248 genomic DNA were assembled using HGAP (hierarchical genome assembly process) using the PacBio Single Molecule Real Time (SMRT) Portal (V2.3.0) (38), with default settings and minimum seed read length of 4,509 bp. Contiguity was used to visualize the assembly and the overlap between contigs using BLASTn (39, 40). The resulting assembly was used as a reference in the RS Resequencing module of PacBio’s SMRT Analysis v2.3.0 to map the raw reads onto the reference genome producing a highly accurate genome consensus. Illumina-sequenced reads of the strain QMA0248 were mapped to the PacBio assembly using Snippy v3.0 (https://github.com/tseemann/snippy). To analyze read pileup in the potential rRNA tandem operon, raw reads of QMA0248 were mapped onto the ~7 Kb rRNA contig using BLASR v2.2, as part of the PacBio’s SMRT Analysis Suite, and visualized using Artemis (41). Methylated DNA bases were identified in the resulting genome assembly using the RS Modification and Motif Analysis protocol and Motif Finder v1 within the SMRT Analysis suite v2.3.0 using Quality Value (QV) cutoff of 30. DNA Methyltransferases (MTases) were identified using nucleotide comparisons against REBASE (42).

### PCR to investigate the rRNA tandem duplication

To investigate the presence of a rRNA tandem repeat, long range PCR from unique adjacent region to unique adjacent region was performed using specific primers for each of 5 rRNA regions (Supplementary Table S6), which were designed using Primer3 (43). PCR was done using LongAmp® Taq PCR Kit (NEB) from 40 ng of QMA0248 *S. iniae* genomic DNA as follows: 5 min at 94°C; 30 cycles of (30 sec at 94°C, 30 sec at 56°C, and 15 min at 65°C); and 10 min final extension at 65°C. The gel was loaded with 5 μL QUICK-LOAD® 1 Kb Extend DNA Ladder and 1μL of PCR products and run for 90 min at 70V using 0.7% TAE buffer solution and stained with HydraGreen.

### Genome and mobile genetic elements annotation

Automated genome annotation was done using Prokka v1.11 (Prokaryotic Genome Annotation System) (44) and then manually curated using Artemis (41) and Geneprimp (45). The start codons of known coding sequences (CDS), such as in the capsule and streptolysin S operons, were further adjusted where appropriate using UniProtKB (http://www.uniprot.org/) and Pfam (46). CRISPRs (Clustered Regularly Interspaced Short Palindromic Repeats) were predicted using CRISPR finder tool (http://crispr.u-psud.fr/crispr/) (47). Prophage annotation was done using PHAST (Phage Search Tool) (http://phast.wishartlab.com/) (48). Island Viewer http://www.pathogenomics.sfu.ca/islandviewer/) was used to predict genomic islands (GI) (49). Boundaries of phage and GI regions were manually adjusted to their respective attachment sites. IS Saga (http://www-genome.biotoul.fr/) was used for the initial identification of insertion sequences (IS). Additional manual curation was carried out to confirm the boundaries of complete and partial IS elements using the IS Finder database (http://www-is.biotoul.fr/). IS element matches against the database that were at ≥95% nucleotide identity were assigned the top matching IS name. IS elements of less percent identity are novel and were assigned the names IS*Stin2*–IS*Stin10* by ISFinder. The impact of IS elements on flanking coding sequences was analyzed using Artemis and by searching the amino acid sequence in UniProt KB (http://www.uniprot.org/), and Pfam databases (41, 46). The complete annotated genome sequence was deposited at GenBank under the accession number CP022392.

### Comparative genomics and phylogenetic analysis

Alignments of the whole-genome or genomic sub-regions, such as the CRISPR and genomic islands, were done using BLASTn implemented in EasyFig v2.1 (40, 50). Detailed analysis of regions of difference and comparison of insertion sequences (IS) were done using Artemis Comparison Tool (ACT) (32). The core genome of the five genomes QMA0248, SF1, YSFST01-82, ISET0901, and ISNO was defined using Roary (51). Phylogenetic trees were constructed using the core genome and the core SNP methods, using multiple programs (see below), using the strain QMA0140 as the out-group (52). The strain *S. iniae* QMA0140 is a dolphin isolate from USA in 1976, and was sequenced using Illumina HiSeq 2000 (See Bioproject PRJNA417543). For quality control, the first 20 bp of each read derived from QMA0140 genomic DNA were hard trimmed using Nesoni v0.132 (https://github.com/Victorian-Bioinformatics-Consortium/nesoni) with a minimum length and quality of 70, and 20, respectively. Hard trimmed filtered reads of QMA0140 were assembled using SPAdes v3.9.0 where contigs < 10X coverage and smaller than 100 bp were removed (53). For core genome phylogenies, whole-genome alignment of the five complete genomes and QMA0140 draft assembly was done using Mauve v2.4.0 and Parsnp v1.2 with default parameter settings (54, 55). For the alignment using Mauve, conserved blocks in all 6 genomes longer than 500 bp were selected and concatenated using the stripSubsetLCBs script producing the core genome alignment. For core SNP phylogenies, error-free simulated reads were created using wgsim v0.3.2 (https://github.com/lh3/wgsim) and mapped to the reference genome QMA0248 along with QMA0140 hard trimmed and filtered raw reads using bowtie v1.0.0 (56), where variants were called using Nesoni v0.132 (https://github.com/Victorian-Bioinformatics-Consortium/nesoni) using default parameters. Core SNPs were also identified by mapping the reads of the 6 genomes to QMA0248 using BWA-MEM v0.7.15 (r1140), implemented in Snippy v3.0 (https://github.com/tseemann/snippy) (57). All trees were produced by RAxML v8.2.9 (58) using the general time-reversible (GTR) and GAMMA distribution model of among-site rate variation with bootstrapping from 1000 replicates, and viewed using FigTree v1.4.0 (http://tree.bio.ed.ac.uk/software/figtree).

## Supplementary Information

### *S. iniae* QMA0248 methylome

DNA methylation guides numerous critical processes including defence against foreign DNA, DNA replication and repair, gene expression and virulence. Analysis of PacBio sequence data enabled the detection of genome-wide DNA methylation. Three DNA methyltransferases (MTases) were encoded in the QMA0248 genome. On the basis of homology to known MTases, QMA0248_0514 (annotated as M.Sin248ORF514P in REBASE) likely targets the GCNGC motif and QMA0248_1949 (annotated as M.Sin248ORF1949P in REBASE) likely targets GCCHR (1). QMA0248_0505 (annotated as M.Sin248ORF0505P in REBASE) is encoded ~5kb upstream of QMA0248_0514 but has no close functional homologs and thus has an unknown recognition sequence (1). A small fraction of GCNGC and GCCHR motifs were methylated in QMA0248 (<3%). Collectively these two motifs account for 21,421 sites across the chromosome (or 1 every 100 bp) suggesting a potential role for methylation in the regulation of gene expression in QMA0248. This figure is roughly equivalent to the frequency of Dam GATC sites in *Escherichia coli*. Further work is required to determine if the activity detected here is biologically meaningful.

### 5-methylcytosine DNA methyltransferase QMA0248_0514

A putative Type II 5-methylcytosine (m5C) DNA MTase (QMA0248_0514, annotated as M.Sin248ORF514P in REBASE) likely targets the GCNGC motif (1). Uncharacterised homologs with 99% amino acid identity are found in 8 other available *S. iniae* complete or draft genomes including SF1, YSFST01-82, ISET0901, ISNO and 89353 (1). In most cases restriction enzymes predicted to recognise GCNGC are predicted to be encoded nearby, or immediately adjacent to the respective MTase gene. Notably, in QMA0248 the adjacent restriction enzyme (QMA0248_0515) is a pseudogene that has been truncated by an IS*981*. In *S. iniae* KCTC 11634BP, the orthologous gene in its draft quality 454 genome was truncated the same point by a contig break (2), suggesting that in both QMA0248 and KCTC 11634BP the MTase does not function as part of a restriction-modification system. The GCNGC motif is found 8074 times in the QMA0248 genome suggesting that methylation activity could have wide-ranging regulatory consequences.

The closest homologs to QMA0248_0514 for which MTase activity has been determined is the M.CmaLM2II enzyme from *Clostridium mangenotii* LM2 and the M.LmoJ3I enzyme from *Listeria monocytogenes* J3115. Despite sharing modest overall amino acid similarity to QMA0248_0514 (59% and 45%, respectively), regions of high amino acid identity within their predicted target recognition domains (34/34 for M.CmaLM2II and 32/24 for M.LmoJ3I) support the prediction that QMA0248_0514 would also methylate the 2^nd^ cytosine of the GCNGC motif. Detection of m5C using PacBio data is normally unreliable (3). As expected, methylation was detected at only a small fraction of GCNGC sites in the QMA0248 genome and the consensus motif determined by the PacBioSMRT-Portal software includes additional bases that are probably artefactual (e.g. GCNGCAGC) (Supplementary Table S5). Further experimental work (such as using Tet1 pre-treatment to enhance detection of m5C with PacBio sequencing, or Oxford Nanopore sequencing) is needed to determine the true extent of cytosine methylation in the *S. iniae* genome and its role in gene regulation.

### N4-methylcytosine DNA methyltransferase QMA0248_1949

The second *S. iniae* QMA0248 MTase (QMA0248_1949, known as M.Sin248ORF1949P in REBASE) shares 95% amino acid identity with the Orphan (gamma) N4-methylcytosine (m4C) DNA MTase M.NgoDCXV (gb|AJ004687.2) from *Neisseria gonorrhoeae*, which specifically targets the GCCHR motif. M.NgoDCXV homologs are remarkably rare and confined to a few streptococcal species (including *S. iniaie* SF1). No specific methylation of GCCHR was detected in the QMA0248 genome but 126 motifs that partially overlapped with GCCHR showed evidence of methylation (Supplementary Table S5). The GCCHR motif is present in 13,347 locations in the QMA0248 genome so this represents only a fraction of available sites. The m4C modification is normally detectable from PacBio sequence data, therefore further work is required to determine if QMA0248_1949 is expressed and functional.

### Putative Type II N4-methylcytosine or N6-methyladenine DNA methyltransferase QMA0248_0505

The third MTase in *S. iniae* QMA0248 (QMA0248_0505, known as M.Sin248ORF0505P in REBASE) is encoded ~5kb upstream of QMA0248_0514 and shares a similar strain distribution. There are no close homologs of QMA0248_0505 for which a recognition site has been determined. Accordingly it has been annotated by REBASE as a putative Type II N4-cytosine or N6-adenine DNA methyltransferase of unknown recognition sequence (1).

## Supplementary Figures

**Figure S1:**
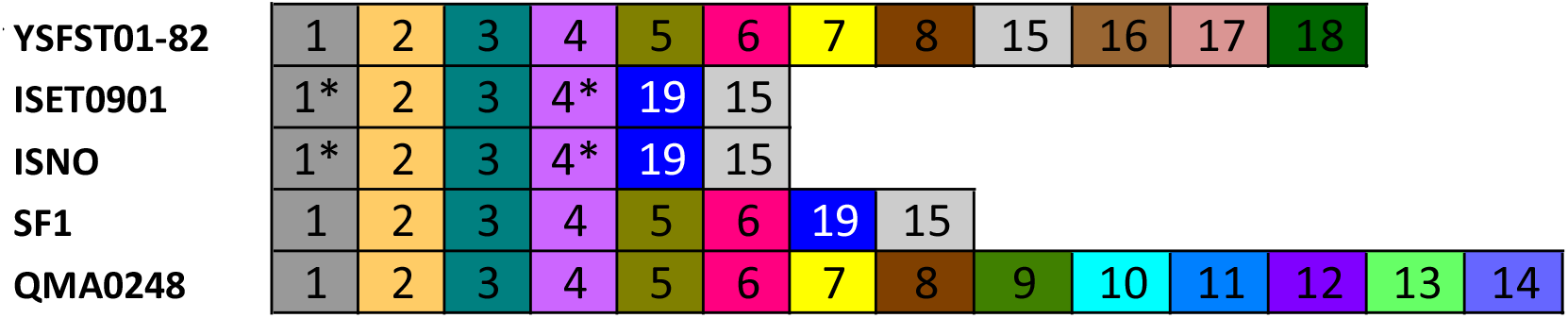
Graphical representation of spacers in CRISPR between YSFST01-82, ISET0901, ISNO, SF1, and QMA0248. Each box represents a spacer, where the same colour and number indicate identical spacers. *: A spacer with an additional direct repeat sequence.

**Figure S2:**
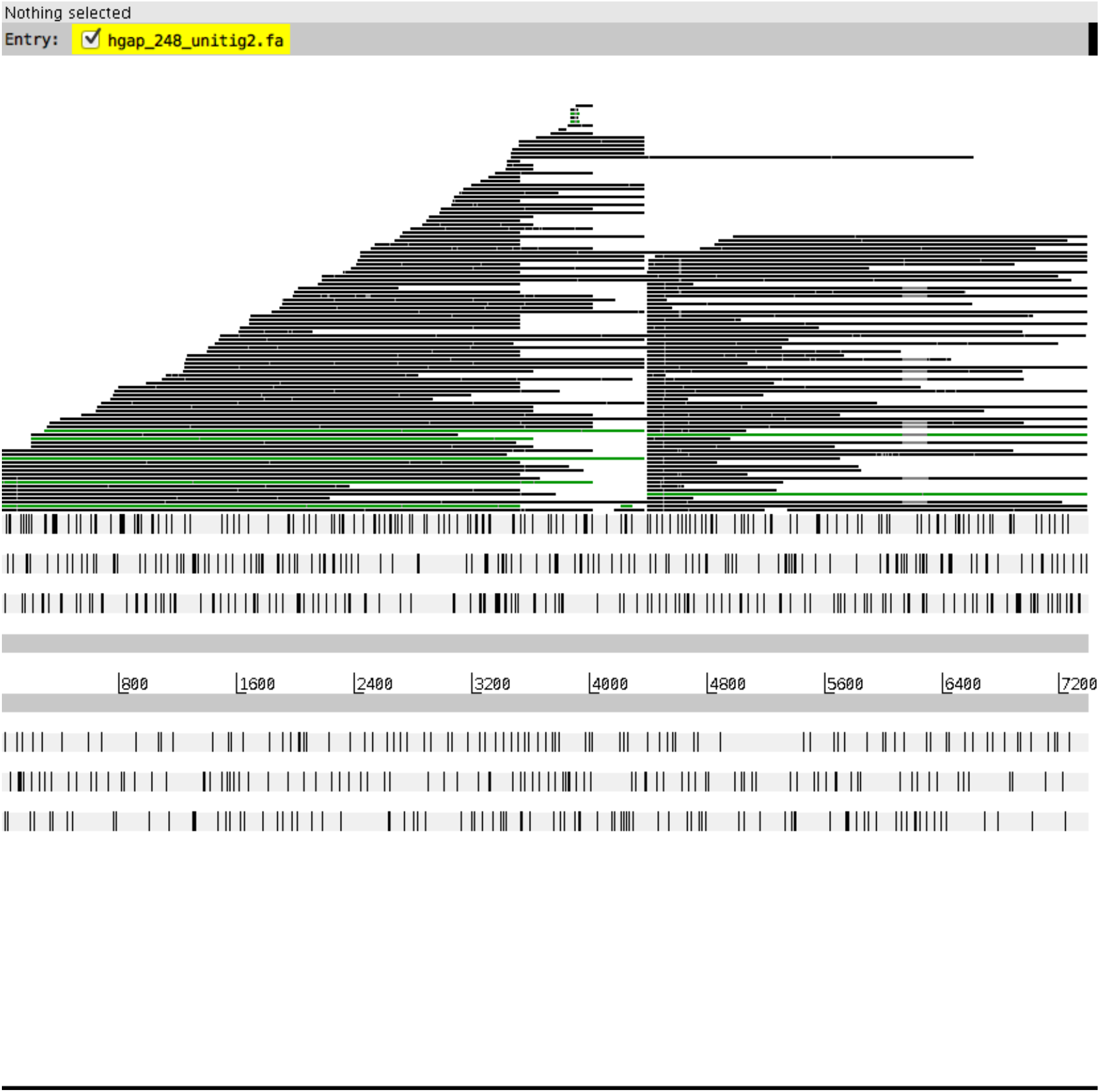
PacBio raw reads pileup of *Streptococcus iniae* QMA0248 on the spurious 7.2 Kb rRNA contig. Alignment of reads is visualised using BamView implemented in Artemis. The lower panel corresponds to the forward and reverse strands of the 7.2 Kb contig, where vertical black lines indicate stop codons. Horizontal lines on the upper panel correspond to the aligned reads, where Artemis colours duplicate reads that span the same region in green black.

**Figure S3:**
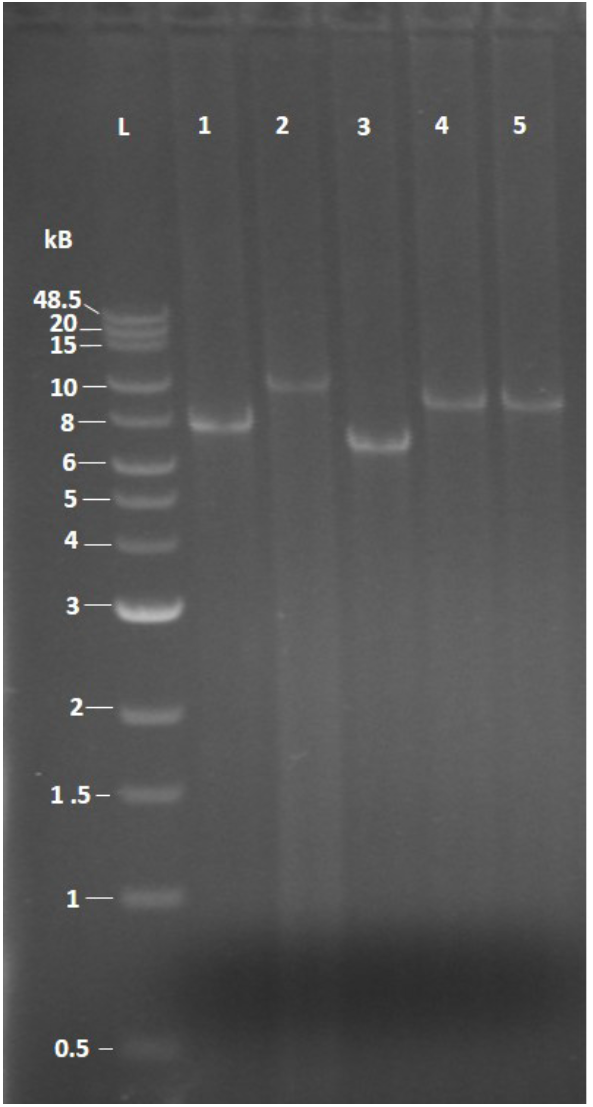
Long-range PCR across rRNA operons. Gel electrophoresis of long range PCR products across five rRNA operons of *S. iniae* QMA0248 (lanes 1-5 in operon-wise order as listed in Table 4). Primers are listed in Table 4. “L” denotes 1 Kb Extend DNA Ladder (NEB).

**Figure S4:**
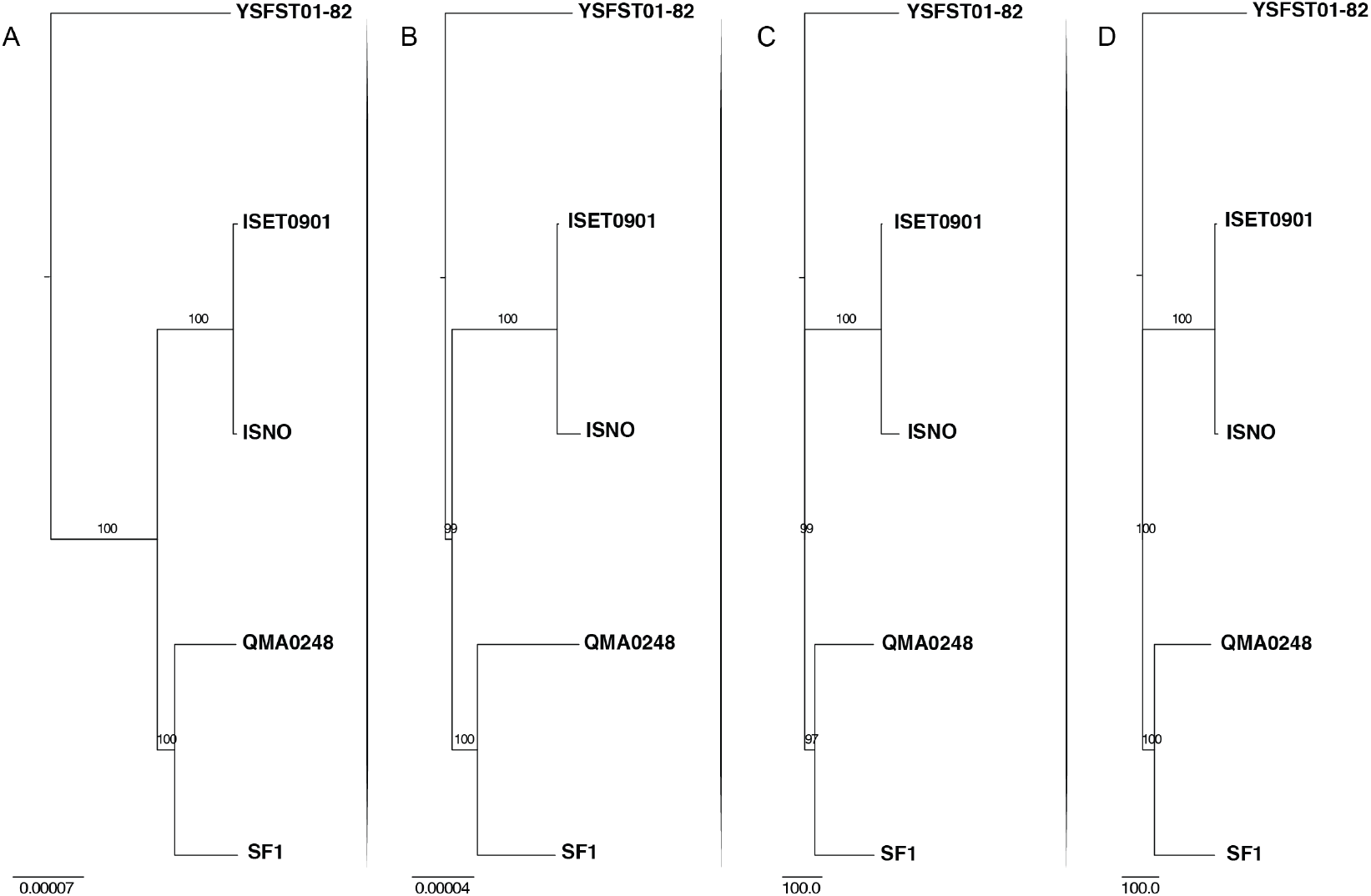
Alternative tree-building methods. Phylogenetic trees constructed for *S. iniae* based on core genome using Mauve (A) and Parsnp (B), and core SNP using Nesoni (C) and Snippy (D). Maximum likelihood (ML) phylograms were built using RAxML where bootstrap support values are shown on branches. Trees are rooted using QMA0140 (not shown). Branch lengths correspond to the number of substitutions (C and D), and the number of substitutions per site (A and B). ML phylograms in A and B were built from 1,928,667 bp and 1,856,820 bp core genome alignment, respectively, whereas C and D were built using 1,241 and 1,116 substitutions, respectively.

## Notes

### Competing Interest Statement

The authors have declared no competing interest.

### Summary of Updates

Supplementary Appendix has been added to PDF. Typographical errors corrected. Discussion updated.

